# BonVision – an open-source software to create and control visual environments

**DOI:** 10.1101/2020.03.09.983775

**Authors:** Gonçalo Lopes, Karolina Farrell, Edward A. B. Horrocks, Chi-Yu Lee, Mai M. Morimoto, Tomaso Muzzu, Amalia Papanikolaou, Fabio R. Rodrigues, Thomas Wheatcroft, Stefano Zucca, Samuel G. Solomon, Aman B. Saleem

**Author notes:** These authors are listed alphabetically (Karolina Farrell, Edward A. B. Horrocks, Chi-Yu Lee, Mai M. Morimoto, Tomaso Muzzu, Amalia Papanikolaou, Fabio R. Rodrigues, Thomas Wheatcroft, Stefano Zucca). These authors jointly supervised this work (Samuel G. Solomon, Aman B. Saleem).

## Abstract

Real-time rendering of closed-loop visual environments is necessary for next-generation understanding of brain function and behaviour, but is prohibitively difficult for non-experts to implement and is limited to few laboratories worldwide. We developed BonVision as an easy-to-use open-source software for the display of virtual or augmented reality, as well as standard visual stimuli. As the architecture is based on the open-source Bonsai graphical programming language, BonVision benefits from native integration with experimental hardware. BonVision therefore enables easy implementation of closed-loop experiments, including real-time interaction with deep neural networks and communication with behavioural and physiological measurement and manipulation devices.

## Introduction

Understanding behaviour and its underlying neural mechanisms requires the ability to construct and control environments that immerse animals and human subjects in complex naturalistic environments that are responsive to their actions. However, most vision research has been performed in non-immersive environments with standard two-dimensional visual stimuli, such as grating patterns or simple figures, using platforms including PsychToolboxy51 or PsychoPy^2,3^. Pioneering efforts to bring gaming-driven advances in computation and rendering have driven the development of immersive closed-loop visual environments^4–6^: STYTRA provides visual stimuli for larval zebrafish in python^7^, ratCAVE is a specialised augmented reality system for rodents in python^5^, FreemoVR provides virtual reality in Ubuntu/Linux^4^, and ViRMEn provides virtual reality in Matlab^6^. But these new platforms are not readily amenable to research paradigms where precise calibration and timing are required. For example, they do not specify an image in egocentric units of visual angle, lack transparent interaction with external hardware, and require advanced programming expertise.

Our initial motivation was to create a visual display software with three key features. First, a integrated, standardised platform that could rapidly switch between traditional visual stimuli (such as grating patterns) and immersive virtual reality. Second, the ability to replicate experimental workflows across different physical configurations (for example, when moving from one to two computer monitors, or moving from flat-screen to spherical projection). Third, the ability for rapid and efficient interfacing with external hardware (needed for experimentation) without development of complex multi-threaded routines. We needed these advances to be provided in an environment such that users with minimal training in programming were able to construct and run complex, closed-loop experimental designs.

We therefore developed BonVision, an open-source software package for the Bonsai graphical programming language^8^, which can generate and display well-defined visual stimuli in 2D and 3D environments. Bonsai is a high-performance event-based language widely used for neuroscience experiments, capable of real-time interfacing with most types of external hardware. BonVision extends Bonsai by providing pre-built GPU shaders and resources for stimuli used in vision research, including movies, along with an accessible, modular interface for composing stimuli and designing experiments. The definition of stimuli in BonVision is independent of the display hardware, allowing for easy replication of workflows across different experimental configurations. Additional unique features include the ability to automatically detect and define the relationship between the observer and the display from a photograph of the experimental apparatus, and to use the outputs of real-time inference methods to determine the position and pose of an observer online, thereby generating augmented reality environments.

## Results

To provide a framework that allowed both traditional visual presentation and immersive virtual reality, we needed to bring these very different ways of defining the visual scene into the same architecture. We achieved this by mapping the 2D retino-centric coordinate frame (i.e. degrees of the visual field) to the surface of a 3D sphere using the Mercator projection (Fig 1A, Suppl. Fig 1). The resulting sphere could therefore be rendered onto displays in the same way as any other 3D environment. We then used “cube mapping” to specify the 360° projection of 3D environments onto arbitrary viewpoints around an experimental observer (human or animal; Fig 1B). Using this process, a display device becomes a window into the virtual environment, where each pixel on the display specifies a vector from the observer through that window. The vector links pixels on the display to pixels in the ‘cube map’, thereby rendering the corresponding portion of the visual field onto the display.

**Figure 1:**
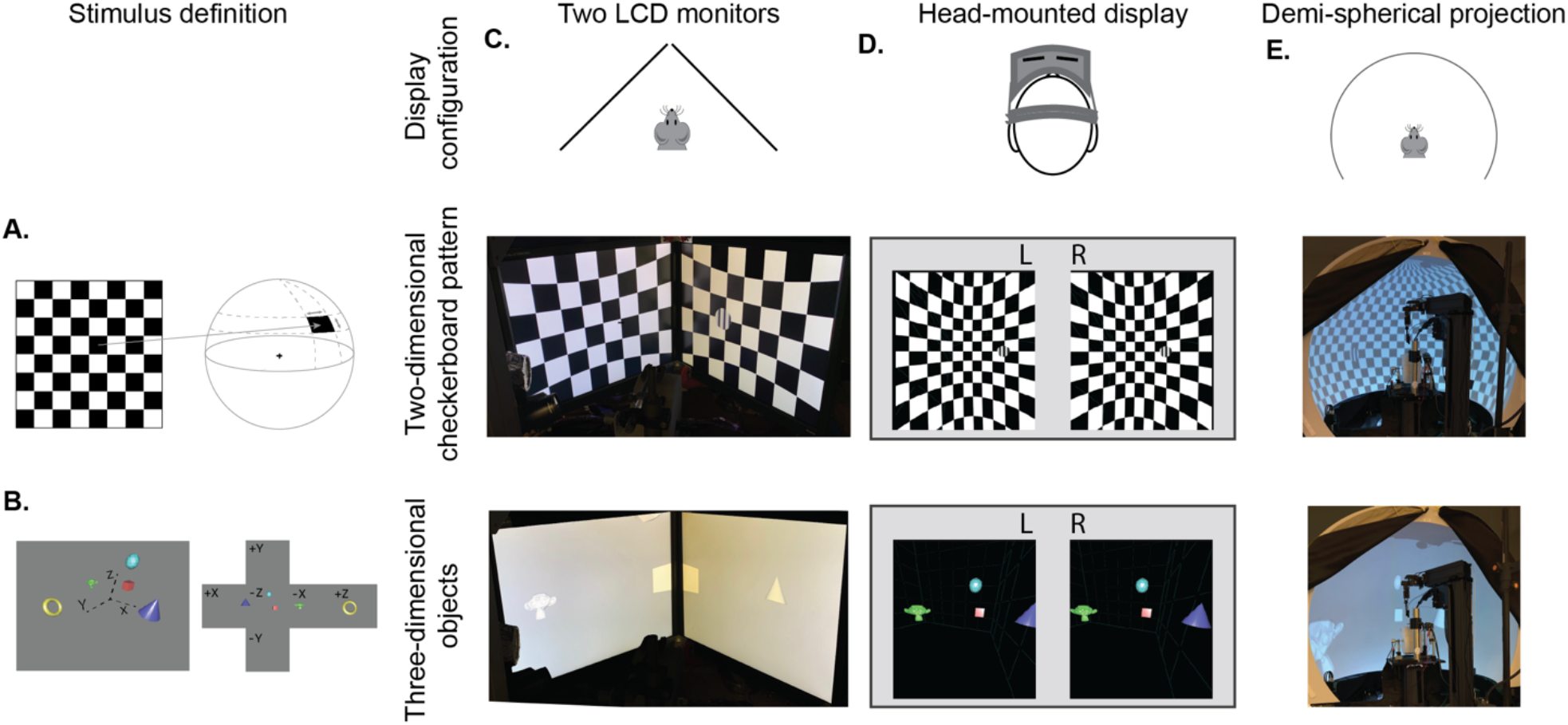
BonVision adaptable display and render configurations. **A.** Illustration of how two-dimensional textures are generated in BonVision using Mercator projection for sphere mapping, with elevation as latitude and azimuth as longitude. **B.** Three-dimensional objects were placed at the appropriate positions and the visual environment was rendered using cube-mapping. **C-E.** Examples of the same two stimuli, a checkerboard + grating (middle row) or four three-dimensional objects (bottom row), displayed in different experimental configurations (top row): two angled LCD monitors (**C**), a head-mounted display (**D**), and demi-spherical dome (**E**).

Our approach has the advantage that the visual stimulus is defined irrespectively of display hardware, allowing us to independently define each experimental apparatus without changing the preceding specification of the visual scene, or the experimental design (Fig 1C-E, Suppl. Fig 1, 2). Consequently, BonVision makes it easy to replicate visual environments and experimental designs on various display devices, including multiple monitors, curved projection surfaces, and head-mounted displays (Fig 1C-E). To facilitate easy and rapid porting between different experimental apparatus, Bonvision features a fast semi-automated display calibration. A photograph of the experimental setup with fiducial markers^9^ measures the 3D position and orientation of each display relative to the observer (Fig 2 and Suppl. Fig. 3). BonVision’s inbuilt image processing algorithms then estimate the position and orientation of each marker to fully specify the display environment.

**Figure 2:**
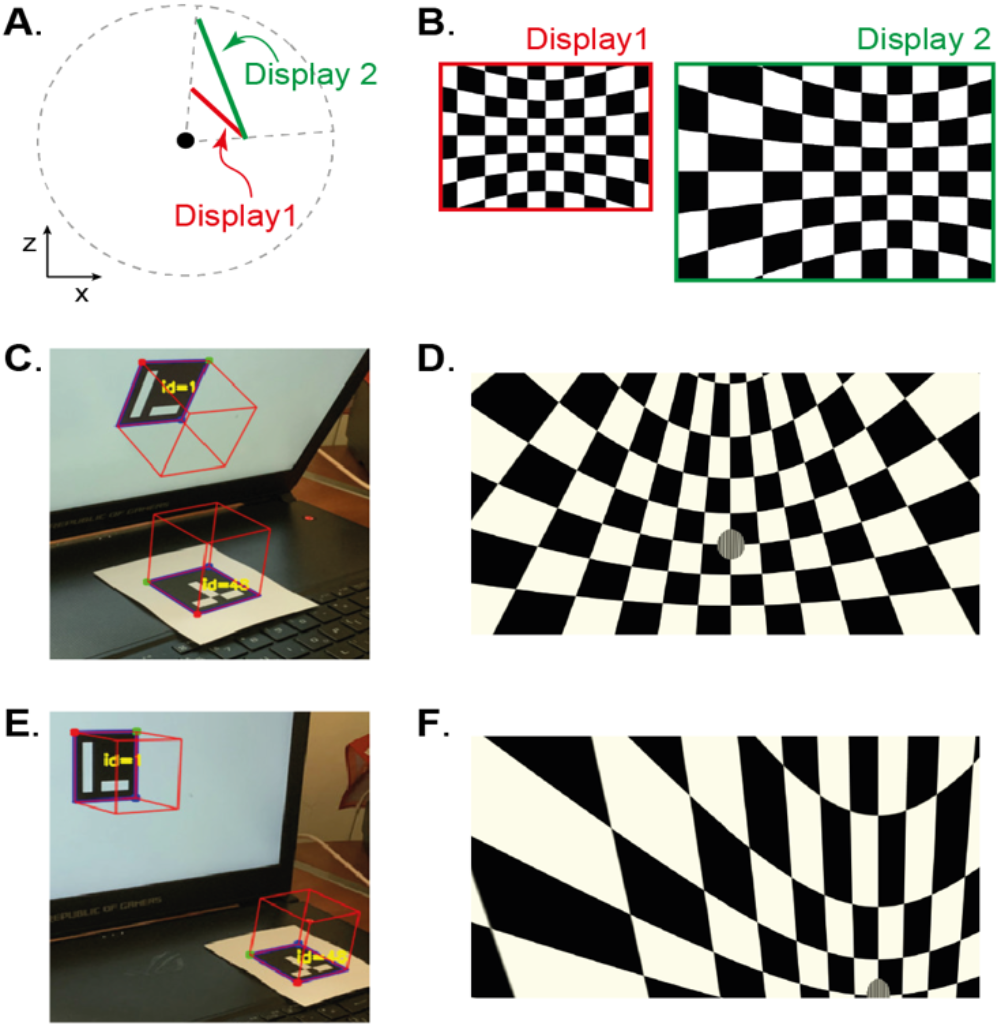
Automated calibration of display position. **A.** Schematic showing the position of two hypothetical displays of different sizes, at different distances and orientation relative to the observer. **B.** How a checkerboard of the same visual angle would appear on each of the two displays. **C.** Example of automatic calibration of display position. Standard markers are presented on the display, or in the environment, to allow automated detection of the position and orientation of both the display and the observer. The superimposed red cubes show the position and orientation of these, as calculated by BonVision. **D.** How the checkerboard would appear on the display when rendered, taking into account the precise position of the display. **E-F.** Same as **C-D,** but for another pair of display and observer positions.

**Figure 3:**
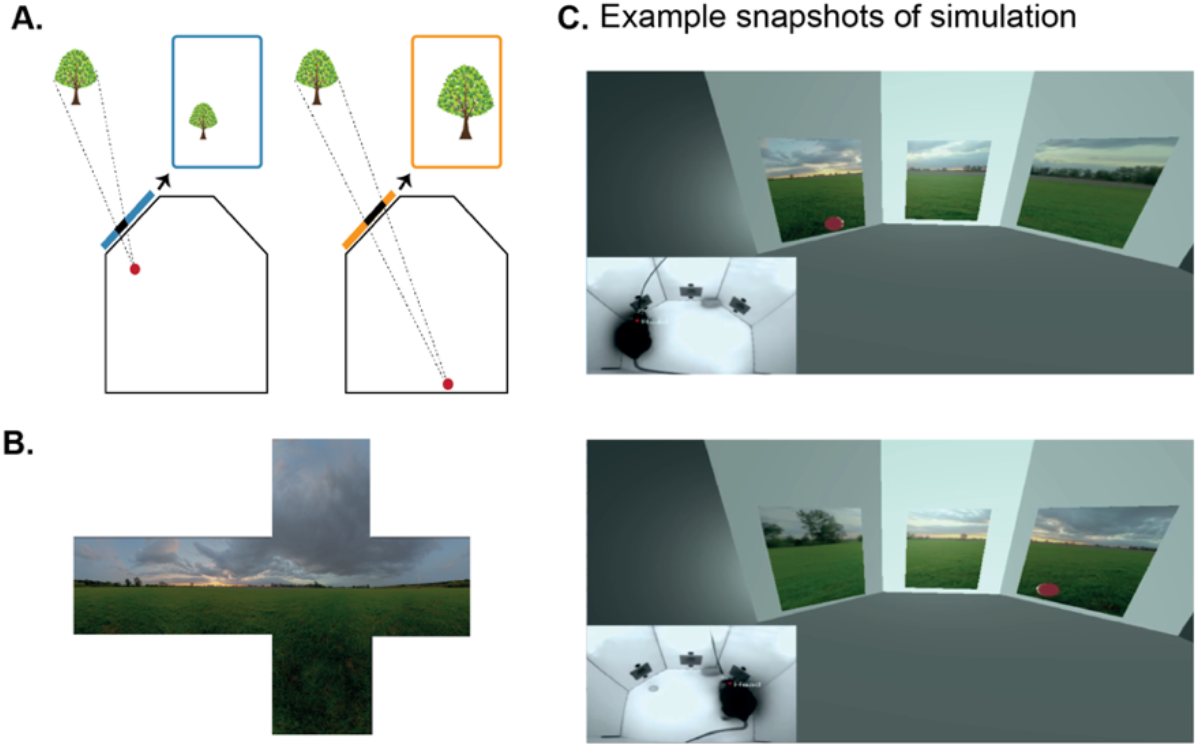
Using BonVision to generate an augmented reality environment. **A.** Illustration of how the image on a fixed display needs to adapt as an animal moves around an environment. The displays simulate windows from a box into a virtual world outside. **B.** The virtual scene (from: http://scmapdb.com/wad:skybox-skies) that was used to generate the example images and Supplementary Video 1. **C.** Two snapshots of scene rendering for instances when the animal was on the left or the right of the environment (from Supplementary Video 1). The inset image shows the video used to determine the viewpoint of the observer: the mouse’s head position was inferred on-line (at a rate of 40 frames/s) by a network trained using DeepLabCut^6^.

Virtual reality environments are easy to generate in BonVision. BonVision has a library of standard pre-defined 3D structures (including planes, spheres and cubes), and environments can be defined by specifying the position and scale of the structures, and the textures rendered on them (e.g. Suppl. Fig. 2 and Fig. 5F). BonVision also has the ability to import standard format 3D design files created elsewhere in order to generate more complex environments. This allows users to leverage existing 3D drawing platforms (including open source platform ‘Blender’) to construct complex virtual scenes.

BonVision can define the relationship between the display and the observer in real-time. This makes it easy to generate augmented reality environments, where what is rendered on a display depends on the position of an observer (Fig 3A). For example, when a mouse navigates through an arena surrounded by displays, BonVision enables closed-loop, position-dependent updating of those displays. BonVision can track markers to determine the position of the observer, but it also has turn-key capacity for real-time live pose estimation techniques – using deep neural networks^10,11^ – to keep track of the observer’s movements. This allows users to generate and present interactive visual environments (Suppl. Video 1, Fig 3B-C).

BonVision is capable of rendering visual environments near the limits of the hardware (Fig 4). This is possible because Bonsai is based on a just-in-time compiler architecture such that there is little computational overhead. To benchmark the responsiveness of BonVision in closed-loop experiments, we measured the delay (latency) between an external event and the presentation of a visual stimulus. We first measured the closed-loop latency for BonVision when a monitor was refreshed at a rate of 60Hz (Fig 4A). We found delays averaged 2.11 ± 0.78 frames (35.26 ± 13.07ms). This latency was slightly shorter than that achieved by PsychToolbox^13^ on the same laptop (2.44 ± 0.59 frames, 40.73 ± 9.8ms; Welch’s t-test, p < 10^−80^, n=1000). The overall latency of BonVision was mainly constrained by the refresh rate of the display device, such that higher frame rate displays yielded lower latency (60Hz: 35.26 ± 13.07ms; 90Hz: 28.45 ± 7.22ms; 144Hz: 18.49 ± 10.1ms; Fig 4A). That is, the number of frames between the external event and stimulus presentation was fairly constant across frame rate (60Hz: 2.11 ± 0.78 frames; 90Hz: 2.56 ± 0.65 frames; 144Hz: 2.66 ± 1.45 frames; Fig 4C). We used two additional methods to benchmark visual display performance relative to other frameworks (we did not try to optimise code fragments for each framework) (Fig 4B-C).

**Figure 4:**
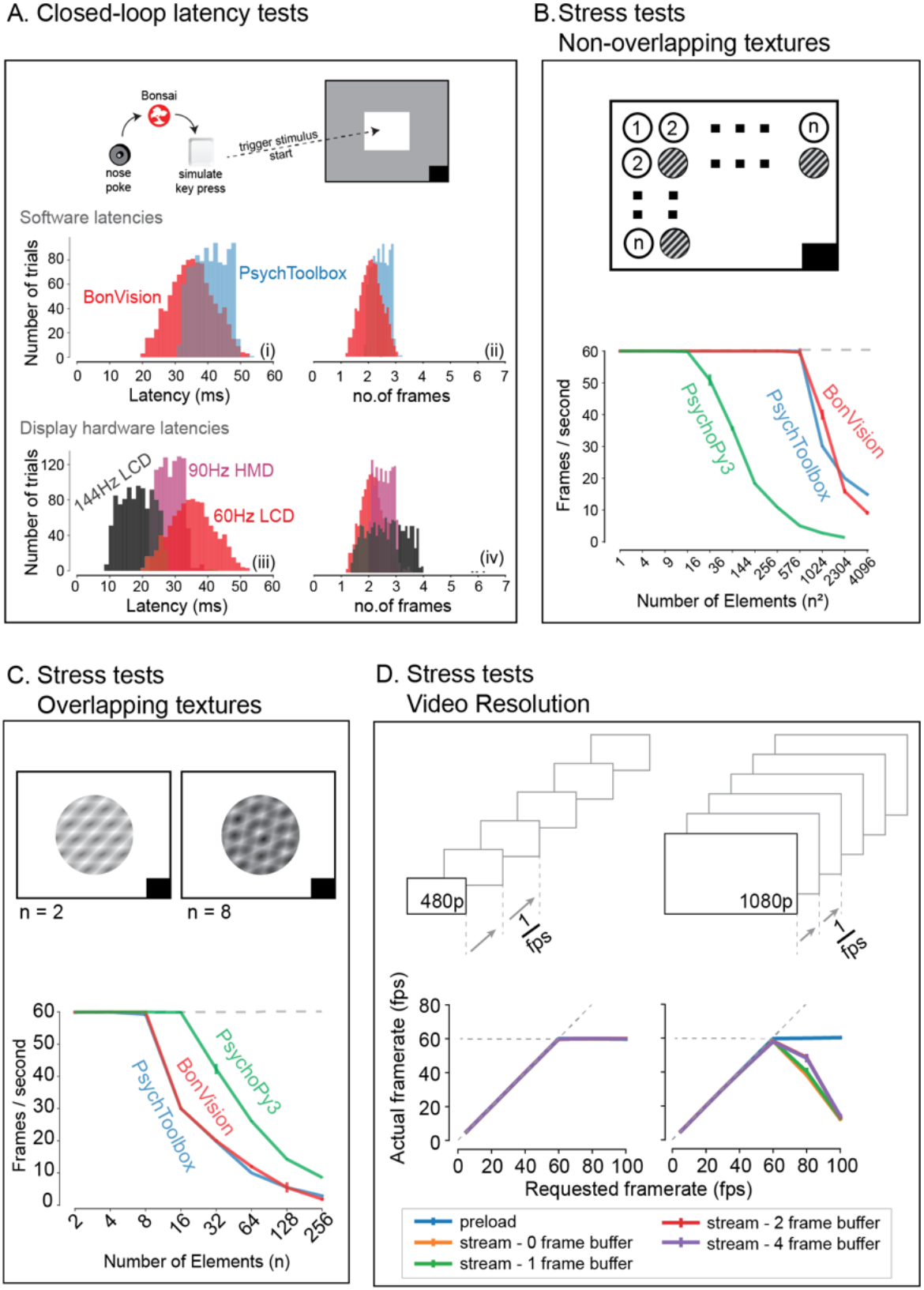
Closed-loop latency and performance benchmarks. **A.** Latency between sending a command (virtual key press) and updating the display (measured using a photodiode). (A.i - A.ii) Latency depended on the frame rate of the display, updating stimuli with a delay of 1-3 frames. (A.iii - A.iv). **B-C.** Benchmarked performance of BonVision with respect to Psychtoolbox and PsychoPy. **B.** When using non-overlapping textures BonVision and Psychtoolbox could present 576 independent textures without dropping frames, while PsychoPy could present 16. **C.** When using overlapping textures PsychoPy could present 16 textures, while BonVision and Psychtoolbox could present 8 textures without dropping frames. **D.** Benchmarks for movie playback. BonVision is capable of displaying standard definition (480p) and high definition (1080p) movies at 60 frames/s on a laptop computer with a standard CPU and graphics card. We measured display rate when fully pre-loading the movie into memory (blue), or when streaming from disk (with no buffer: orange; 1-frame buffer: green; 2-frame buffer: red; 4-frame buffer: purple). When asked to display at rates higher than the monitor refresh rate (>60 frames/s), the 480p video played at the maximum frame rate of 60fps in all conditions, while the 1080p video reached the maximum rate when pre-loaded. Using a buffer slightly improved performance.

BonVision was able to render up to 576 independent elements and up to 8 overlapping textures at 60Hz without missing (‘dropping’) frames, broadly matching PsychoPy^2,3^ and Psychtoolbox^1^. BonVision also supports video playback, either by preloading the video or by streaming it from the disk. The streaming mode, which utilises real-time file I/O and decompression, is capable of displaying both standard definition (SD: 480p) and full HD (HD: 1080p) at 60Hz on a standard computer (Fig 4D). At higher rates, performance is impaired for Full HD videos, but is improved by buffering, and fully restored by preloading the video onto memory (Fig 4D). We benchmarked BonVision on a standard Windows OS laptop, but BonVision is now also capable of running on Linux.

To confirm that the rendering speed and timing accuracy of BonVision is sufficient to support neurophysiological experiments, which need high timing accuracy, we mapped the receptive fields of neurons early in the visual pathway^12^, in the primary visual cortex and superior colliculus. The stimulus (‘sparse noise’) consisted of small black or white squares briefly (0.1s) presented at random locations (Fig 5A). This stimulus, which is commonly used to measure receptive fields of visual neurons, is sensitive to the timing accuracy of the visual stimulus, meaning that errors in timing would prevent the identification of receptive fields. In our experiments using BonVision, we were able to recover receptive fields from electrophysiological measurements^13^, both in the superior colliculus and primary visual cortex of awake mice (Fig 5B-C), demonstrating that BonVision meets the timing requirements for visual neurophysiology.

**Figure 5:**
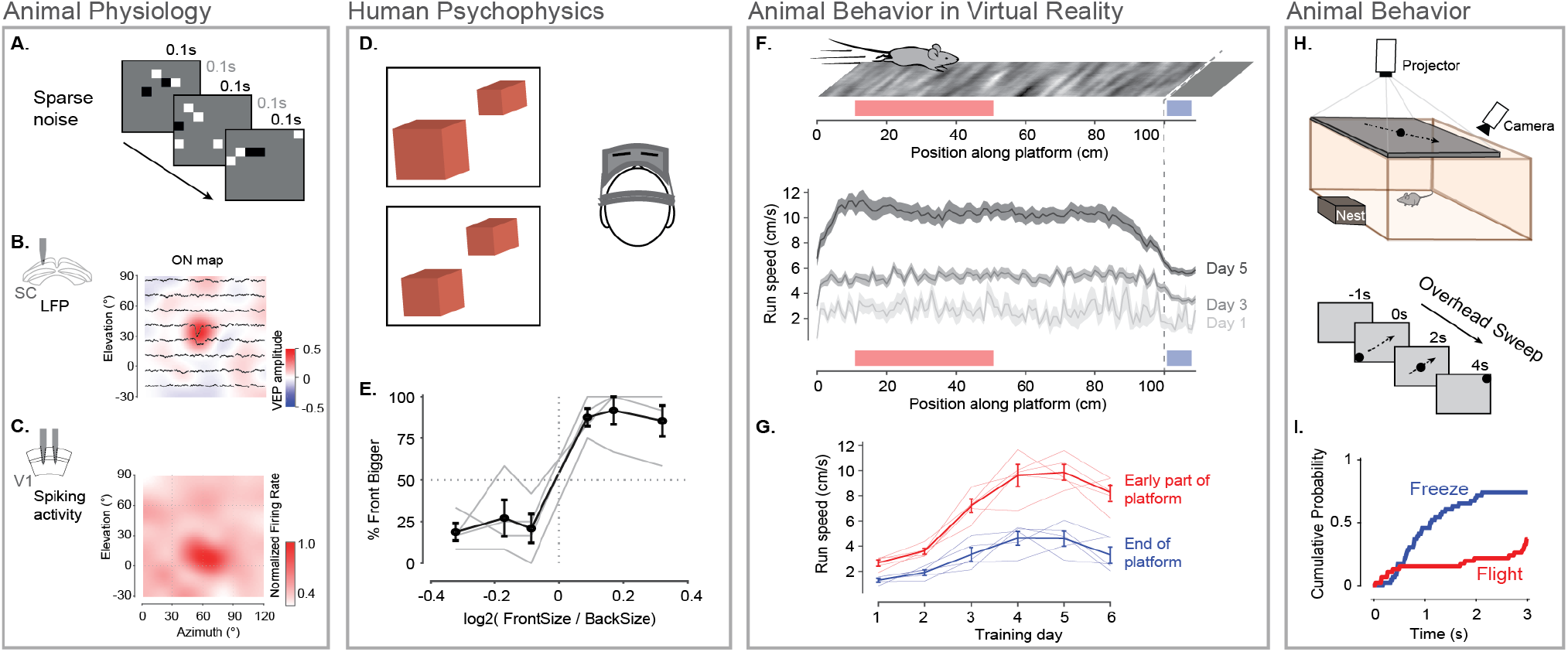
Illustration of BonVision across a range of vision research experiments. **A.** Sparse noise stimulus, generated with BonVision, is rendered onto a demi-spherical screen. **B-C**. Receptive field maps from recordings of local field potential in the superior colliculus (**B**), and spiking activity in the primary visual cortex (**C**) of mouse. **D.** Two cubes were presented at different depths in a virtual environment through a head-mounted display to human subjects. Subjects had to report which cube was larger: left or right. **E.** Subjects predominantly reported the larger object correctly, with a slight bias to report that the object in front was bigger. **F.** BonVision was used to generate a closed-loop virtual platform that a mouse could explore (top: schematic of platform). Mice naturally tended to run faster along the platform, and in later sessions developed a speed profile, where they slowed down as they approached the end of the platform (virtual cliff). **G.** The speed of the animal at the start of the platform and at the end of the platform as a function training. **H.** BonVision was used to present visual stimuli overhead while an animal was free to explore an environment (which included a refuge). The stimulus was a small dot (5° diameter) moving across the projected surface over several seconds. **I.** The speed of the animal across different trials, aligned to the time of stimulus appearance.

To assess whether BonVision could create immersive virtual reality environments we first asked human observers to discriminate the size of objects presented at different depths on a head-mounted display^14^. BonVision uses positional information (obtained from the head-mounted display) to update the view of the world that needs to be provided to each eye, and returns two appropriately rendered images. On each trial, the observer identified the larger of two non-overlapping cubes that were placed at different virtual depths (Fig 5D-E). The display was updated in closed-loop to allow observers to alter their viewpoint by moving their head. Distinguishing objects of the same retinal size required observers to use depth-dependent cues^15^, and we found that all observers were able to identify which cube was larger (Fig 5E). We next projected a simple environment onto a dome that surrounded a head-fixed mouse (as shown in Fig 1E). The mouse was free to run on a treadmill, and the treadmill’s movements were used to update the mouse’s position on a virtual platform (Fig 5F). Mouse locomotion speed increased with repeated exposure (Fig 5F-G), suggesting habituation to the virtual environment. Strikingly, mouse speed varied with position on the virtual platform (Fig 5F-G), reducing rapidly at the end of the platform, where the environment provided a virtual cliff. This suggests that BonVision is capable of eliciting naturalistic behaviours in a virtual environment. BonVision was also able to produce instinctive avoidance behaviours in freely-moving mice (Fig 5H-I). We displayed a small black dot slowly sweeping across the overhead visual field. Visual stimuli presented in BonVision primarily elicited a freezing response, which similar experiments have previously described^10^ (Fig 5I). Together these results show that BonVision provides sufficient rendering performance to support human and animal visual behaviour.

## Discussion

BonVision is a single software package to support experimental designs that require visual display, including virtual and augmented reality environments. BonVision is easy and fast to implement, cross-platform and open source, providing versatility and reproducibility.

BonVision addresses several persistent barriers to reproducibility in visual experiments. First, BonVision is able to reproduce and deliver visual stimuli on very different experimental apparatus. This is possible because BonVision’s architecture separates specification of the display and the visual environment. Second, BonVision includes a library of workflows and operators to standardize and ease the construction of new stimuli and virtual environments. For example, it has established protocols for defining display positions (Suppl. Fig 3), mesh-mapping of curved displays (Fig 1E), and automatic linearization of display luminance (Suppl. Fig 4), as well as a library of examples for experiments commonly used in visual neuroscience. In addition, the modular structure of BonVision enables the development and exchange of custom nodes for generating new visual stimuli or functionality without the need to construct the complete experimental paradigm. Third, BonVision is based on Bonsai^8^, which has a large user base and an active developer community, and is now a standard tool for open-source neuroscience research. BonVision naturally integrates Bonsai’s established packages in the multiple domains important for modern neuroscience, which are widely used in applications including real-time video processing^16,17^, optogenetics^16–18^, fibre photometry^19,20^, electrophysiology (including specific packages for Open Ephys^13,21^ and high-density silicon probes^22,23^), and calcium imaging (e.g. UCLA miniscope^24,25^). Bonsai is specifically designed for flexible and high-performance composition of data streams and external events, and is able to monitor and interconnect multiple sensor and effector systems in parallel, thus making it particularly easy to implement closed-loop experiments.

In summary, BonVision can generate complex 3D environments and retinotopically defined 2D visual stimuli within the same framework. Existing platforms used for vision research, including PsychToolbox^1^, PsychoPy^2,3^, STYTRA^7^, or RigBox^26^, focus on well-defined 2D stimuli. Similarly, gaming-driven software, including FreemoVR^4^, ratCAVE^5^, and ViRMEn^6^, are oriented towards generating virtual reality environments. BonVision combines the advantages of both these approaches in a single framework (Supplementary Table 1), while bringing the unique capacity to automatically calibrate the display environment, and use deep neural networks to provide real-time control of virtual environments. Experiments in BonVision can be rapidly prototyped and easily replicated across different display configurations. Being free, open-source and portable, BonVision is a state-of-the-art tool for visual display that is accessible to the wider community.

## Code availability

BonVision is an open-source software package available to use under the MIT license. It can be downloaded through the Bonsai (bonsai-rx.org) package manager, and the source code is available at: github.com/bonvision/BonVision. Installation instructions, demos and learning tools are available at: bonvision.github.io/.

## Acknowledgements

We are profoundly thankful to Bruno Cruz and Joe Paton for sharing their videos of mouse behaviour. This work was supported by a Wellcome Enrichment award: Open Research (200501/Z/16/A), Sir Henry Dale Fellowship from the Wellcome Trust and Royal Society (200501), Human Science Frontiers Program grant (RGY0076/2018) to A.B.S., an International Collaboration Award (with Adam Kohn) from the Stavros Niarchos Foundation / Research to Prevent Blindness to S.G.S., Medical Research Council grant (R023808), Biotechnology and Biological Sciences Research Council grant (R004765) to S.G.S. and A.B.S.

## Author Contributions

This work was conceptualised by G.L., S.G.S. and A.B.S., the software was developed by G.L., methodology and validation were by all authors, writing – original draft was by G.L., S.G.S. and A.B.S., and writing – review & editing was by G.L., K.F., M.M.M., T.M., F.R.R., T.W., S.Z., S.G.S. & A.B.S, and supervision and funding acquisition was by S.G.S. and A.B.S.

---

**Supplementary Table 1:** Features of visual display software

| Features | BonVision | PsychToolBox | PsychoPy | ViRMEn | ratCAVE | FreemoVR | Unity |
| --- | --- | --- | --- | --- | --- | --- | --- |
| Free and Open-Source (FOSS) | ✓✓ | ✓# | ✓✓ | ✓# | ✓ | ✓✓ | ✓ |
| Rendering of 3D environments | ✓✓ | ✓ | ✓ | ✓✓ | ✓✓ | ✓✓ | ✓✓ |
| Dynamic rendering based on observer viewpoint | ✓✓ |  |  | ✓ | ✓✓ | ✓✓ | ✓ |
| Import 3<sup>rd</sup> party 3D scenes | ✓✓ | ✓ | ✓ |  |  |  | ✓✓ |
| Interfacing with cameras, sensors, and effectors | ✓✓ | ✓✓ | ~ | ✓✓ |  | ~ | ~ |
| Real-time hardware control | ✓✓ | ~ | ~ | ✓ | ✓✓ | ✓ | ✓ |
| Traditional visual stimuli | ✓✓ | ✓✓ | ✓✓ |  |  |  |  |
| Auto-calibration of display position and pose | ✓✓ |  |  |  |  |  |  |
| Integration with deep learning pose estimation | ✓✓ |  |  |  |  |  |  |

- ✓✓ easy and well-supported ✓ possible or not well-supported ~ difficult to implement

**Supplementary Video 1: Augmented reality using BonVision.** This video is an example of a deep neural network, trained with DeepLabCut, being used to estimate the position of a mouse’s head in an environment in real-time, and updating a virtual scene presented on the monitors based on this estimated position. The first few seconds of the video display the online tracking of specific features (nose, head, and base of tail) while an animal is moving around (shown as a red dot) in a three-port box (Soares, Atallah & Paton, 2016). The inset shows the original video the simulation is based on. The rest of the video demonstrates how a green field landscape (source: http://scmapdb.com/wad:skybox-skies) outside the box is displayed on the three displays within the box. The three displays simulate windows into the world beyond the box. The position of the animal was updated at 40 frames/s.

## Supplementary Figures

**Supplementary Fig 1. (related to Figures 1 and 2):**
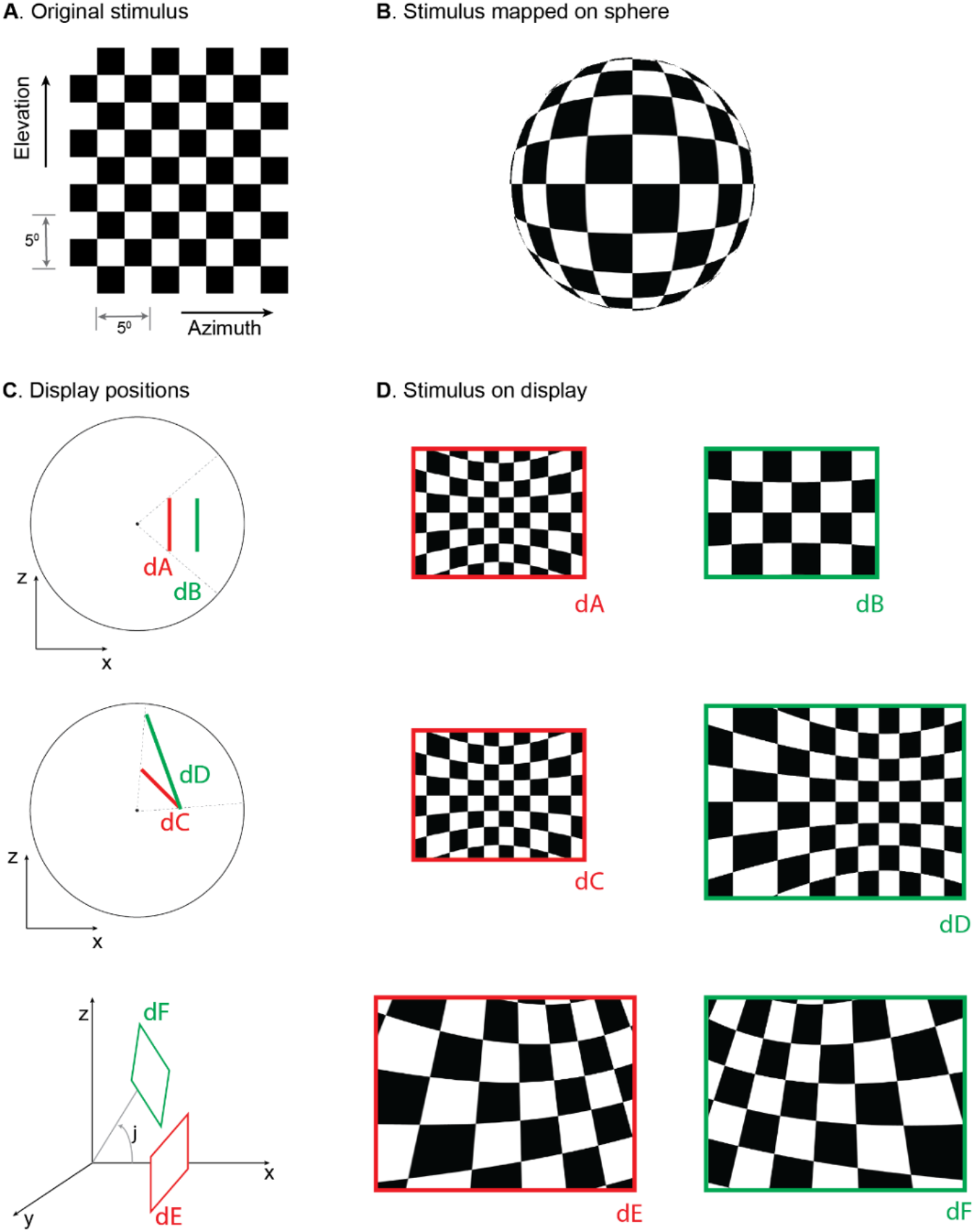
Mapping stimuli onto displays in various positions. **A.** Checkerboard stimulus being rendered. **B.** Projection of the stimulus onto a sphere using Mercator projection. **C.** Example display positions (dA-dF) and (**D**) corresponding rendered images.

**Supplementary Fig 2. (related to Figure 1):**
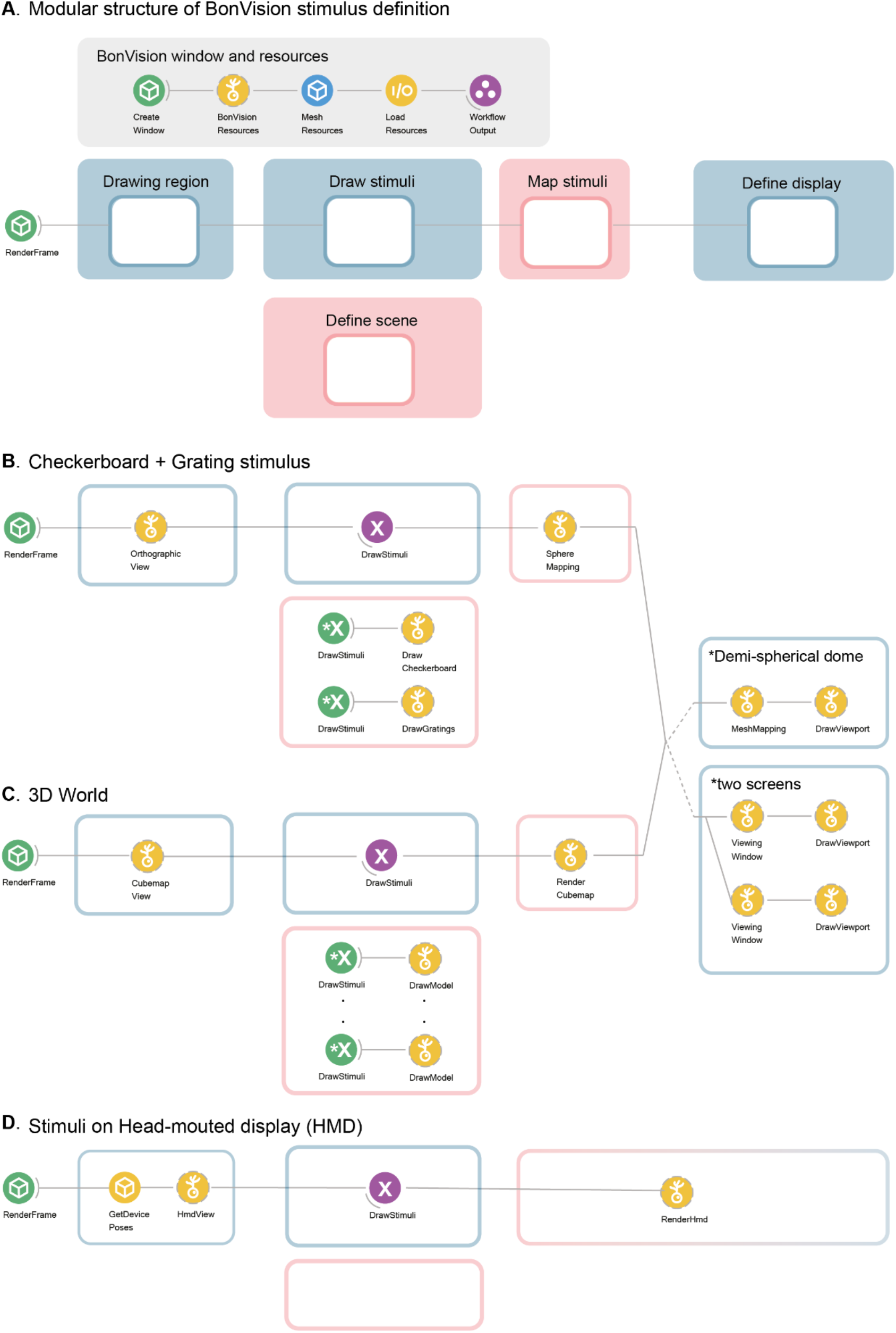
Modular structure of workflow and example workflows. **A.** Description of the modules in BonVision workflows that generate stimuli. Every BonVision stimuli includes a module that creates and initializes the render window, shown in “BonVision window and resources”. This defines the window parameters in *Create Window* (such as background colour, screen index, VSync), and loads predefined (*BonVision Resources*) and user defined textures (*Texture Resources*, not shown), and 3D meshes (*Mesh Resources*). This is followed by the modules: “Drawing region”, where the visual space covered by the stimuli is defined, which can be the complete visual space, 360° x 360°. “Draw stimuli” and “Define scene” are where the stimulus is defined, “Map Stimuli”, which maps the stimuli into the 3D environment, and “Define display”, where the display devices are defined. **B-C**. Modules that define the checkerboard + grating stimulus (**B**) shown in the middle row of Fig 1, and 3D world (**C**) with 5 objects shown in the bottom row of Fig 1. The display device is defined separately and either display can be appended at the end of the workflow. This separation of the display device allows for replication between experimental configurations. **D**. The variants of the modules used to display stimuli on a head-mounted display. The empty region under “Define scene” would be filled by the corresponding nodes in **B** and **C**.

**Supplementary Figure 3. (related to Figure 2):**
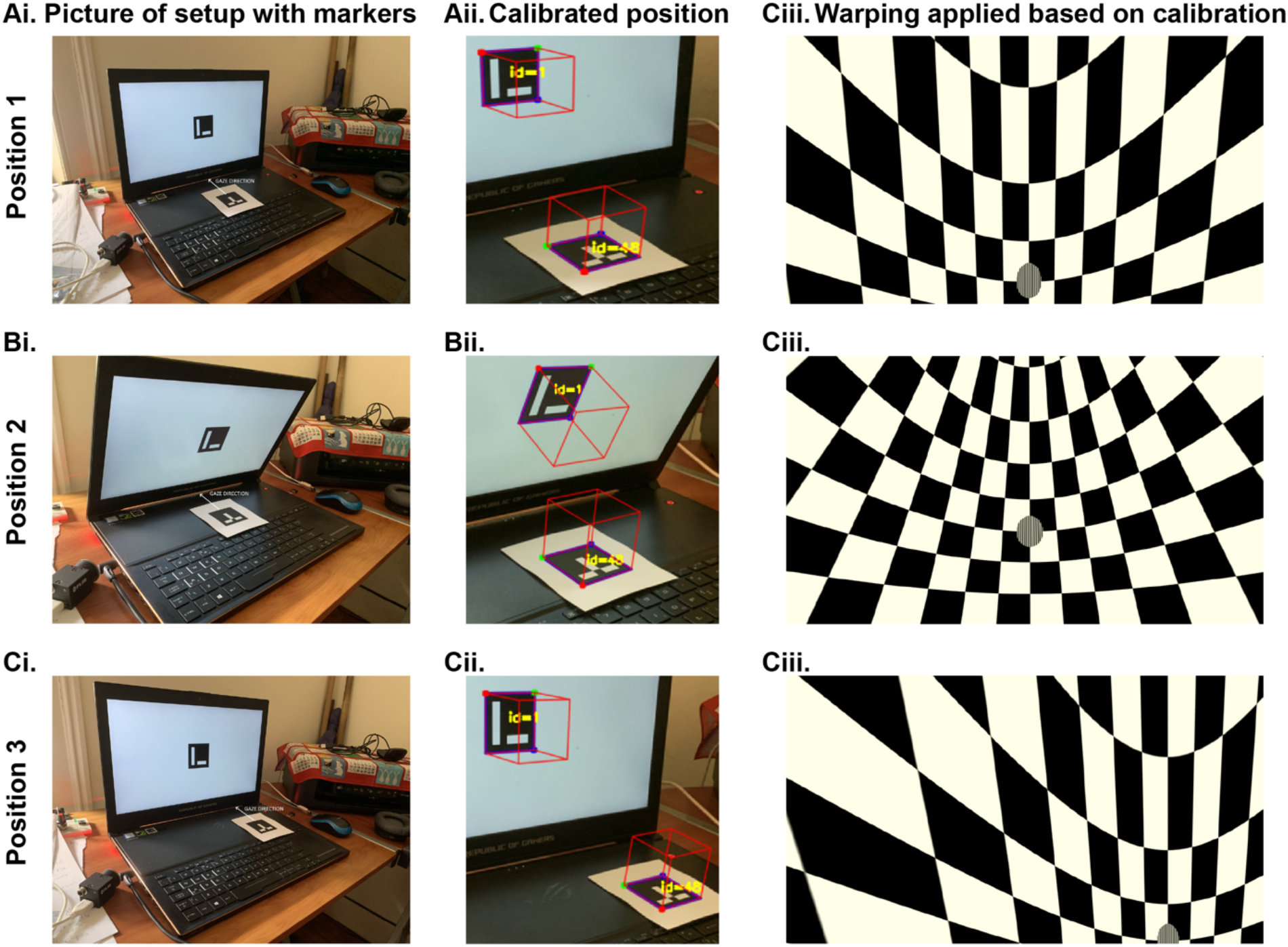
Automated workflow to calibrate display position. The automated calibration is carried out by taking advantage of ArUco markers^5^ that can be used to calculate the 3D position of a surface. **Ai.** We use one marker on the display and one placed in the position of the observer. We then use a picture of the display and observer position taken by a calibrated camera. This is an example where we used a mobile phone camera for calibration. **Aii.** The detected 3D positions of the screen and the observer, as calculated by BonVision. **Aiii.** A checkerboard image rendered based on the precise position of the display. **B-C**. same as A-C for different screen and observer positions: with the screen tilted towards the animal (**B**), or the observer shifted to the right of the screen (**C**).

**Supplementary Figure 4:**
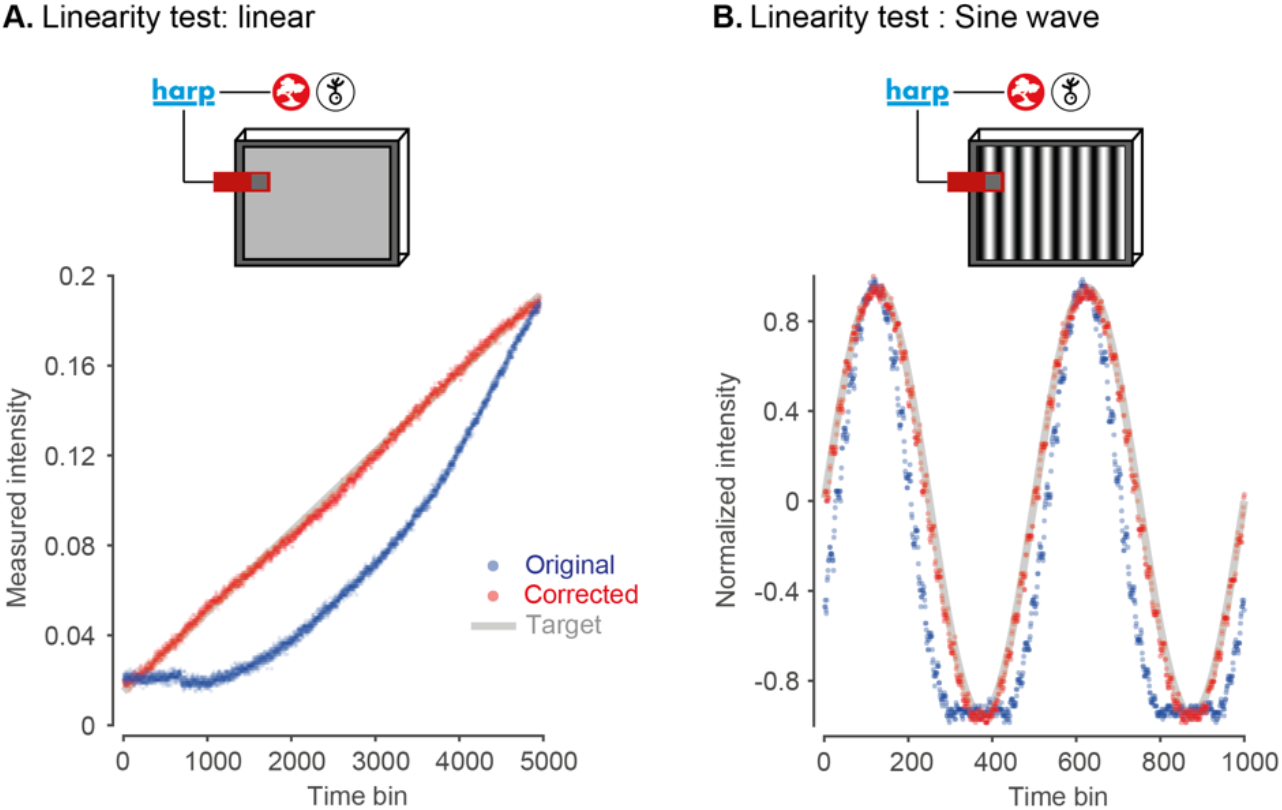
Automated gamma-calibration of visual displays. BonVision monitored a photodiode (Photodiode v2.1, https://www.cf-hw.org/harp/behavior) through a HARP microprocessor, to measure the light output of the monitor (Dell Latitude 7480). The red, green and blue channels of the display were sent the same values (i.e. grey scale). **A.** Gamma calibration. The input to the display channels was modulated by a linear ramp (range 0-255). Without calibration the monitor output (arbitrary units) increased exponentially (blue line). The measurement was then used to construct an intermediate look-up table that corrected the values sent to the display. Following calibration, the display intensity is close to linear (red line). Inset at top: schematic of the experimental configuration. **B.** Similar to A, but showing the intensity profile of a drifting sinusoidal grating. Measurements before calibration resemble an exponentiated sinusoid (blue dotted line). Measurements after calibration resemble a regular sinusoid (red dotted line).

## Experimental Procedures

### Benchmarking

We performed benchmarking to measure latencies and skipped (“dropped”) frames. For benchmarks at 60Hz refresh rate, we used a standard laptop with the following configuration: Dell Latitude 7480, Intel Core i7-6600U Processor Base with Integrated HD Graphics 520 (Dual Core, 2.6GHz), 16GB RAM. For higher refresh rates we used a gaming laptop ASUS ROG Zephyrus GX501GI, with an Intel Core i7-8750H (6 cores, 2.20GHz), 16GB RAM, equipped with a NVIDIA GeForce GTX 1080. The gaming laptop built-in display refreshes at 144Hz, and for measuring latencies at 90Hz we connected it to a Vive Pro SteamVR head-mounted display (90Hz refresh rate). All tests were run on Windows 10 Pro 64-bit.

To measure the time from input detection to display update, as well as dropped frames detection, we used open-source HARP devices from Champalimaud Research Scientific Hardware Platform. Specifically we used the HARP Behavior device (https://www.cf-hw.org/harp/behavior) to synchronise all measurements with the extensions: ‘Photodiode v2.1’ to measure the change of the stimulus on the screen, and ‘Mice poke simple v1.2’ as the nose poke device to externally trigger changes. To filter out the infrared noise generated from an internal LED sensor inside the Vive Pro HMD, we positioned an infrared cut-off filter between the internal headset optics and the photodiode. Benchmarks for video playback were carried out using a trailer from the Durian Open Movie Project (© copyright Blender Foundation | durian.blender.org).

All benchmark programs and data are available at https://github.com/bonvision/benchmarks.

### File Formats

We tested the display of images and videos using the image and video benchmark workflows. We confirmed the ability to use the following image formats: PNG, JPG, BMP, TIFF, GIF. Movie display relies on the FFmpeg library (https://ffmpeg.org/), an industry standard, and we confirmed ability to use the following containers: AVI, MP4, OGG, OGV and WMV; in conjunction with standard codecs: H264, MPEG4, MPEG2, DIVX. Importing 3D models and complex scenes relies on the Open Asset Importer Library (Assimp | http://assimp.org/). We confirmed the ability to import and render 3D models and scenes from the following formats: OBJ, Blender.

### Animal Experiments

All experiments were performed in accordance with the Animals (Scientific Procedures) Act 1986 (United Kingdom) and Home Office (United Kingdom) approved project and personal licenses. The experiments were approved by the University College London Animal Welfare Ethical Review Board under Project License 70/8637. The mice (C57BL6 wild-type) were group-housed with a maximum of five to a cage, under a 12-hour light/dark cycle. All behavioural and electrophysiological recordings were carried out during the dark phase of the cycle.

### Innate Defensive Behaviour

Mice (5 male, C57BL6, 8 weeks old) were placed in a 40cm square arena. A dark refuge placed outside the arena could be accessed through a 10cm door in one wall. A DLP projector (Optoma GT760) illuminated a screen 35cm above the arena with a grey background (80 candela/m^2^). When the mouse was near the centre of the arena, a 2.5cm black dot appeared on one side of the projection screen and translated smoothly to the opposite side over 3.3s. 10 trials were conducted over 5 days and the animal was allowed to explore the environment for 5-10 minutes before the onset of each trial.

Mouse movements were recorded with a near infrared camera (Blackfly S, BFS-U3-13Y3M-C, sampling rate: 60Hz) positioned over the arena. An infrared LED was used to align video and stimulus. Freezing was defined as a drop in the animal speed below 2cm/s that lasted more than 0.1s; flight responses as an increase in the animal running speed above 40cm/s. Responses were only considered if they occurred within 3.5s from stimulus onset.

### Surgery

Mice were implanted with a custom-built stainless-steel metal plate on the skull under isoflurane anaesthesia. A ~1mm craniotomy was performed either over the primary visual cortex (2mm lateral and 0.5mm anterior from lambda) or superior colliculus (0.5mm lateral and 0.2mm anterior from lambda). Mice were allowed to recover for 4-24 hours before the first recording session.

We used a virtual reality apparatus similar to those used in previous studies (Schmidt-Hieber & Hausser, 2013; Muzzu, Mitolo, Gava & Schultz, 2018). Briefly, mice were head-fixed above a polystyrene wheel with a radius of 10cm. Mice were positioned in the geometric centre of a truncated spherical screen onto which we projected the visual stimulus. The visual stimulus was centred at +60° azimuth and +30° elevation and had a span of 120° azimuth and 120° elevation.

### Virtual reality behaviour

5 male, 8-week old, C57BL6 mice were used for this experiment. One week after the surgery, mice were placed on a treadmill and habituated to the Virtual Reality (VR) environment by progressively increasing the number of time spent head fixed: from ~15 mins to 2 hours. Mice spontaneously ran on the treadmill, moving through the VR in absence of reward. The VR environment was a 100cm long platform with a patterned texture that animals ran over for multiple trials. Each trial started with an animal at the start of the platform and ended when it reached the end, or if 60s had elapsed. At the end of a trial, there was a 2 second grey interval before the start of the next trial.

### Neural Recordings

To record neural activity, we used multi-electrode array probes with two shanks and 32 channels (ASSY-37 E-1, Cambridge Neurotech Ltd., Cambridge, UK). Electrophysiology data was acquired with an OpenEphys acquisition board connected to a different computer from that used to generate the visual stimulus.

The electrophysiological data from each session was processed using Kilosort 1 (Pachitariu, Steinmetz, Kadir, Carandini & Harris, 2016). We synchronised spike times with behavioural data by aligning the signal of a photodiode that detected the visual stimuli transitions (PDA25K2, Thorlabs, Inc., USA). We sampled the firing rate at 60Hz, and then smoothed it with a 300ms Gaussian filter. We calculated receptive fields as the average firing rate or local field potential elicited by the appearance of a stimulus in each location (custom routines in MATLAB).

### Augmented reality for mice

The mouse behaviour videos were acquired by Bruno Cruz from the lab of Joe Paton at the Champalimaud Centre for the Unknown, using methods similar to Soares, Atallah & Paton, 2016. A *ResNet-50* network was trained using DeepLabCut (Mathis et al, 2018). We simulated a visual environment in which a virtual scene was presented beyond the arena, and updated the scenes on three walls of the arena that simulated how the view of these objects changed as the animal moved through the environment. The position of the animal was updated from the video file at a rate of 40 frames/s on a gaming laptop: ASUS ROG Zephyrus GX501GI, with an Intel Core i7-8750H (6 cores, 2.20GHz), 16GB RAM, equipped with a NVIDIA GeForce GTX 1080, using a 512×512 video. The performance can be improved using a lower pixel resolution for video capture, and we were able to achieve up to 80 frames/s without noticeable decrease in tracking accuracy using this strategy. Further enhancements can be achieved using a *MobileNet* network. The position inference from the deep neural network and the BonVision visual stimulus rendering were run on the same machine.

### Human Psychophysics

All procedures were approved by the Experimental Psychology Ethics Committee at University College London. 4 male participants were tested for this experiment. The experiments were run on a gaming laptop (described above) connected it to a Vive Pro SteamVR head-mounted display (90Hz refresh rate). BonVision is compatible with different headsets (for example Oculus Rift, HTC Vive). BonVision receives the projection matrix (perspective projection of world display) and the view matrix (position of eye in the world) for each eye from the head set. BonVision uses these matrices to generate two textures, one for the left eye and one for the right eye. Standard onboard computations on the headset provide additional non-linear transformations that account for the relationship between the eye and the display (such as lens distortion effects).

